## Supplementary figures and images for "Microglial trogocytosis and the complement system regulate axonal pruning *in vivo*"

### Supplemental Figure 1

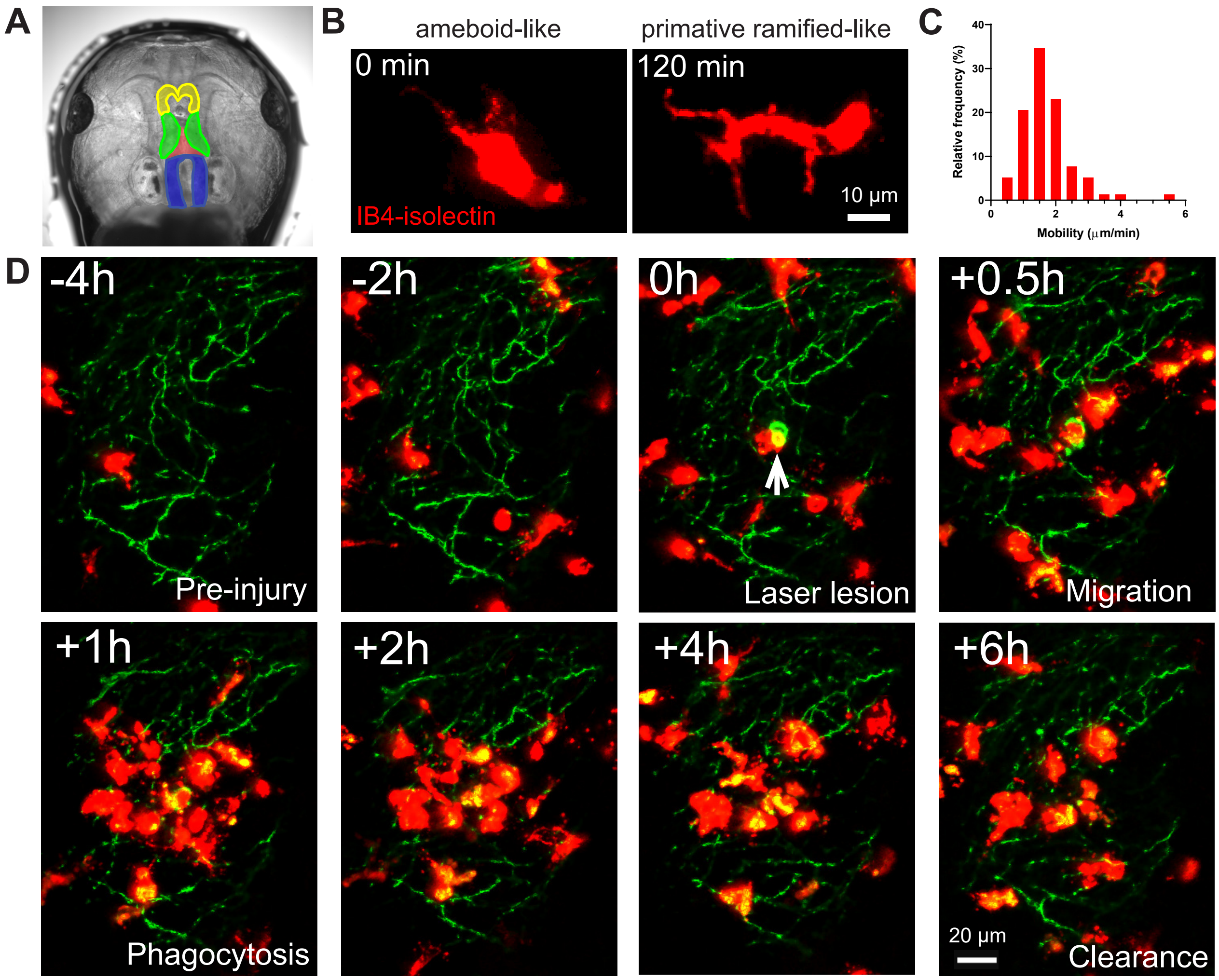

### Supplemental Figure 2

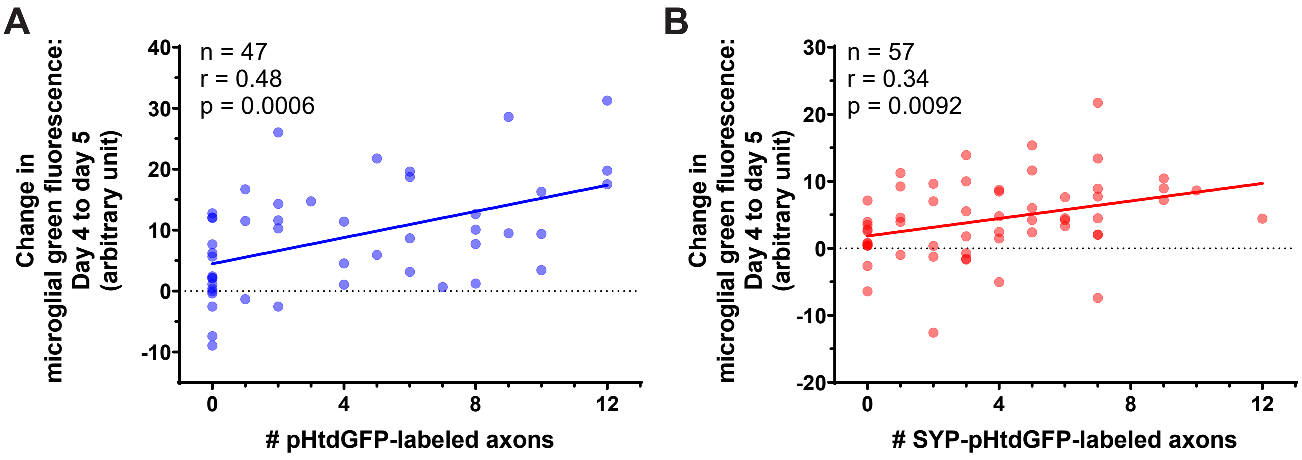

### Supplemental Figure 3

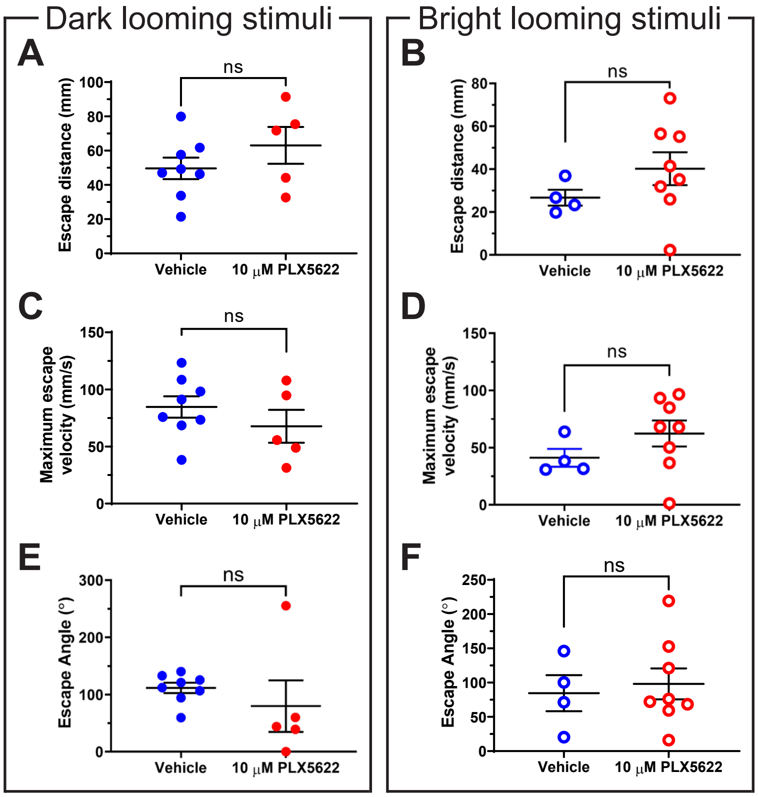

### Supplemental Figure 4

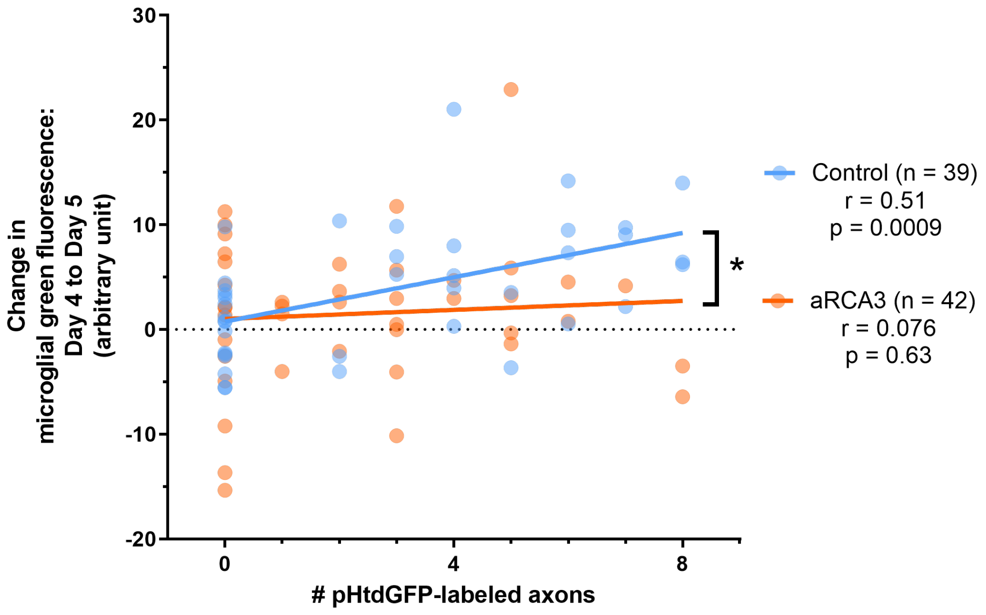
